# Reduced acetylation of TFAM promotes bioenergetic dysfunction in the failing heart

**DOI:** 10.1101/2022.10.31.514513

**Authors:** Manling Zhang, Ning Feng, Dharendra Thapa, Michael W. Stoner, Janet R. Manning, Charles F. McTiernan, Xue Yang, Zishan Peng, Michael J. Jurczak, Danielle Guimaraes, Sruti Shiva, Brett A. Kaufman, Michael N. Sack, Iain Scott

## Abstract

General Control of Amino-Acid Synthesis 5-like 1 (GCN5L1) was previously identified as a key regulator of protein lysine acetylation in mitochondria. Subsequent studies demonstrated that GCN5L1 regulates the acetylation status and activity of mitochondrial fuel substrate metabolism enzymes. However, the role of GCN5L1 in response to chronic hemodynamic stress is largely unknown. Here, we show that cardiomyocyte-specific GCN5L1 knockout mice (cGCN5L1 KO) display exacerbated pressure overload-induced heart failure progression following transaortic constriction (TAC). Mitochondrial DNA and mitochondrial electron transport chain protein levels were decreased in cGCN5L1 KO hearts after TAC, and isolated neonatal cardiomyocytes with reduced GCN5L1 expression had lower bioenergetic output in response to hypertrophic stress. Loss of GCN5L1 expression led to a decrease in the acetylation status of mitochondrial transcription factor A (TFAM) after TAC *in vivo*, which was linked to a reduction in mtDNA levels *in vitro*. Together, these data suggest that GCN5L1 may protect from hemodynamic stress by maintaining mitochondrial bioenergetic output.

**Highlights:**

- Reduced GCN5L1 expression in the failing heart promotes contractile dysfunction
- Mitochondrial DNA (mtDNA) levels are reduced in cardiomyocyte-specific GCN5L1 knockout mice following hemodynamic stress
- GCN5L1 knockdown reduces, and GCN5L1 overexpression increases, bioenergetic output in hypertrophic cardiomyocytes
- GCN5L1-mediated acetylation of TFAM promotes increased mtDNA levels

## Introduction

Heart failure remains one of the leading causes of death and hospital admission in the US (Benjamin et al., 2018). Despite improved therapeutic approaches the prognosis of heart failure remains poor, with ~50% mortality in 5 years (Yancy et al., 2017; Yancy et al., 2013). As such, there is considerable enthusiasm for further exploration of the pathogenic mechanisms underlying heart failure, and the development of novel mechanism-based therapeutic strategies. Metabolic derangements and poor cardiac energy production are hallmarks of heart failure (Bayeva et al., 2013; Bhatt and Butler, 2018; Doehner et al., 2014), and systems that regulate cardiac bioenergetics represent a potential target for therapeutic intervention. Mitochondrial bioenergetic output is controlled by several key mechanisms related to fuel import, substrate oxidation, and electron transport complex function. Mitochondrial Transcription Factor A (TFAM) controls the replication, copy number, and stability of mitochondrial DNA (mtDNA), and is necessary for the maintenance of the mitochondrial electron transport chain (Taherzadeh-Fard et al., 2011; Ventura-Clapier et al., 2008). The abundance of mtDNA is decreased in heart failure, and overexpression of TFAM improves cardiac function after myocardial infarction (Ikeuchi et al., 2005). To date, the molecular mechanisms governing the regulation of TFAM in response to pathological stresses in the heart remain poorly understood.

General Control of Amino-Acid Synthesis 5-like 1 (GCN5L1) was previously identified as a mitochondrial acetyltransferase-related protein (Scott et al., 2012). GCN5L1-mediated acetylation is implicated in the regulation of fatty acid oxidation, glucose oxidation, and mitochondrial respiration under various conditions (Thapa et al., 2020; Thapa et al., 2018; Thapa et al., 2017). However, whether GCN5L1 plays a role in the regulation of cardiac bioenergetics in response to hemodynamic stress is unknown. Using novel cardiac-specific GCN5L1 KO mice (cGCN5L1 KO), we show that GCN5L1 is required to protect hearts against pressure overload-induced heart failure. Mechanistic studies reveal that GCN5L1 enhances cardiac energetics in response to pressure overload (transaortic constriction [TAC]) by maintaining mtDNA abundance via lysine acetylation of TFAM. These data suggest that approaches which support GCN5L1 expression and/or TFAM acetylation may offer therapeutic benefits in future translational studies on heart failure.

## Results

### Cardiomyocyte-specific GCN5L1 KO mice display exacerbated heart failure after TAC

To investigate the role of GCN5L1 in heart failure, we first examined the expression of GCN5L1 in human and mouse failing hearts. We found that GCN5L1 mRNA expression was significantly reduced in explanted hearts from patients with non-ischemic cardiomyopathy, and in mouse hearts subjected to TAC (Fig 1A). The reduction in GCN5L1 mRNA in mouse failing hearts was matched by a decrease in GCN5L1 protein abundance in the same tissues (Fig 1B). To determine the importance of decreased GCN5L1 levels in heart failure development, cardiac-specific GCN5LI knockout (cGCN5L1 KO) mice were generated by crossing floxed GCN5L1 mice with constitutive α-myosin heavy chain (MHC)-Cre mice (Fig S1). Cardiac function in cGCN5L1 KO mice was similar to WT mice to at least one year of age under nonstressed conditions (Fig S1). To understand the role of GCN5L1 during cardiac stress, wildtype (WT) littermate controls and cGCN5L1 KO mice aged 10-12 weeks were subjected to TAC. Compared to WT littermate controls, cGCN5L1 KO mice developed accelerated heart failure progression after TAC, with increased dilation of left ventricular chamber (Fig 1C,D), decreased fractional shortening (Fig 1E), increased heart weight (Fig 1F), and augmented fetal gene (Fig 1G-H) and fibrosis marker (Fig 1I) expression. Male and female mouse data was combined based on the observation that male and female cGCN5L1 KO mice showed similar outcomes after TAC compared to WT controls (Fig S2). Overall, these studies suggest that cardiac GCN5L1 expression is required to protect the heart against pathological stress.

**Figure 1.**
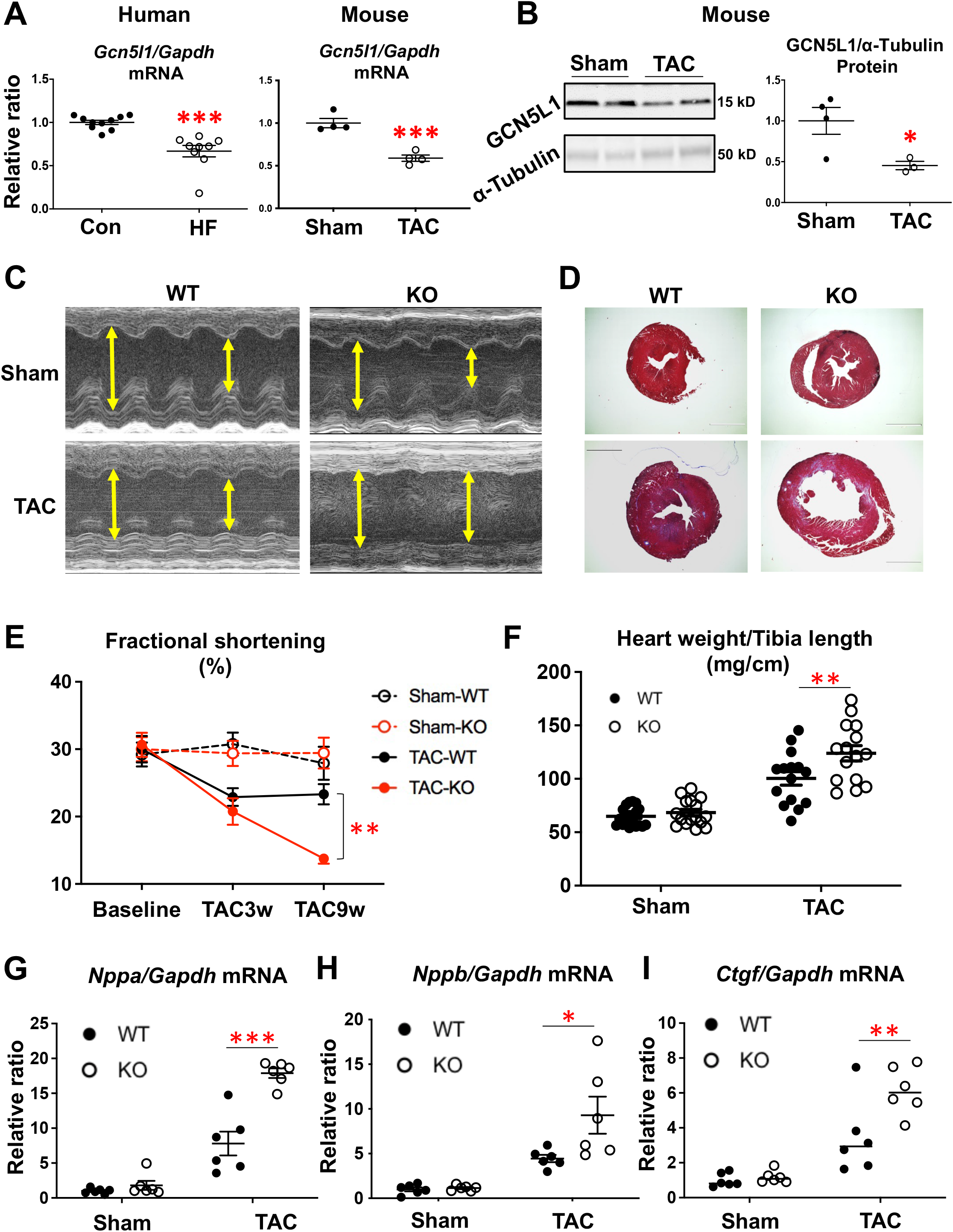
Cardiac-specific GCN5L1 KO mice display exacerbated heart failure in response to pressure overload. (A) GCN5L1 mRNA level was significantly decreased in patients with non-ischemic cardiomyopathy (n = 9-10) and mouse failing hearts after TAC (n = 4). (B) GCN5L1 protein level was significantly decreased in mouse failing hearts (n = 3-4). (CD) Representative echocardiography and histology of WT and cGCN5L1 KO mice subjected to sham or TAC. Scale bar = 1 mm. (E) Compared to WT mice and MHC-Cre controls, cGCN5L1 KO mice showed further decreased fractional shortening after TAC (n = 15). (F) cGCN5L1 KO mouse heart weight was increased relative to WT controls after TAC (n = 15). (G-I) ANP (*Nppa*), BNP (*Nppb*) and CTGF (*Ctgf*) mRNA were increased in cGCN5L1 KO mice after TAC (n = 6). Data are shown as means ± SEM. *P<0.05, **P<0.01, ***P<0.001 vs. TAC-WT group.

### Ablation of GCN5L1 expression in cardiomyocytes reduces ETC protein abundance, mtDNA levels, and bioenergetic output

We next examined the mechanism underlying decreased cardiac function in cGCN5L1 KO mice in response to pressure overload. Strikingly, the abundance of mitochondrial electron transport chain (ETC) proteins from Complex I, III, IV, and the ATP synthase were significantly reduced in cGCN5L1 KO mice after TAC relative to WT controls (Fig 2A). In contrast, there was no difference in protein abundance from Complex II (Fig 2A), which unlike the other complexes contains no mtDNA gene products. We therefore examined mtDNA levels in mice after sham or TAC surgery, and found that cGCN5L1 KO mice displayed a significant decrease in mt-ND2 (complex I) and mt-ATP6 (ATP synthase) expression after TAC relative to their WT controls (Fig 2B).

**Figure 2.**
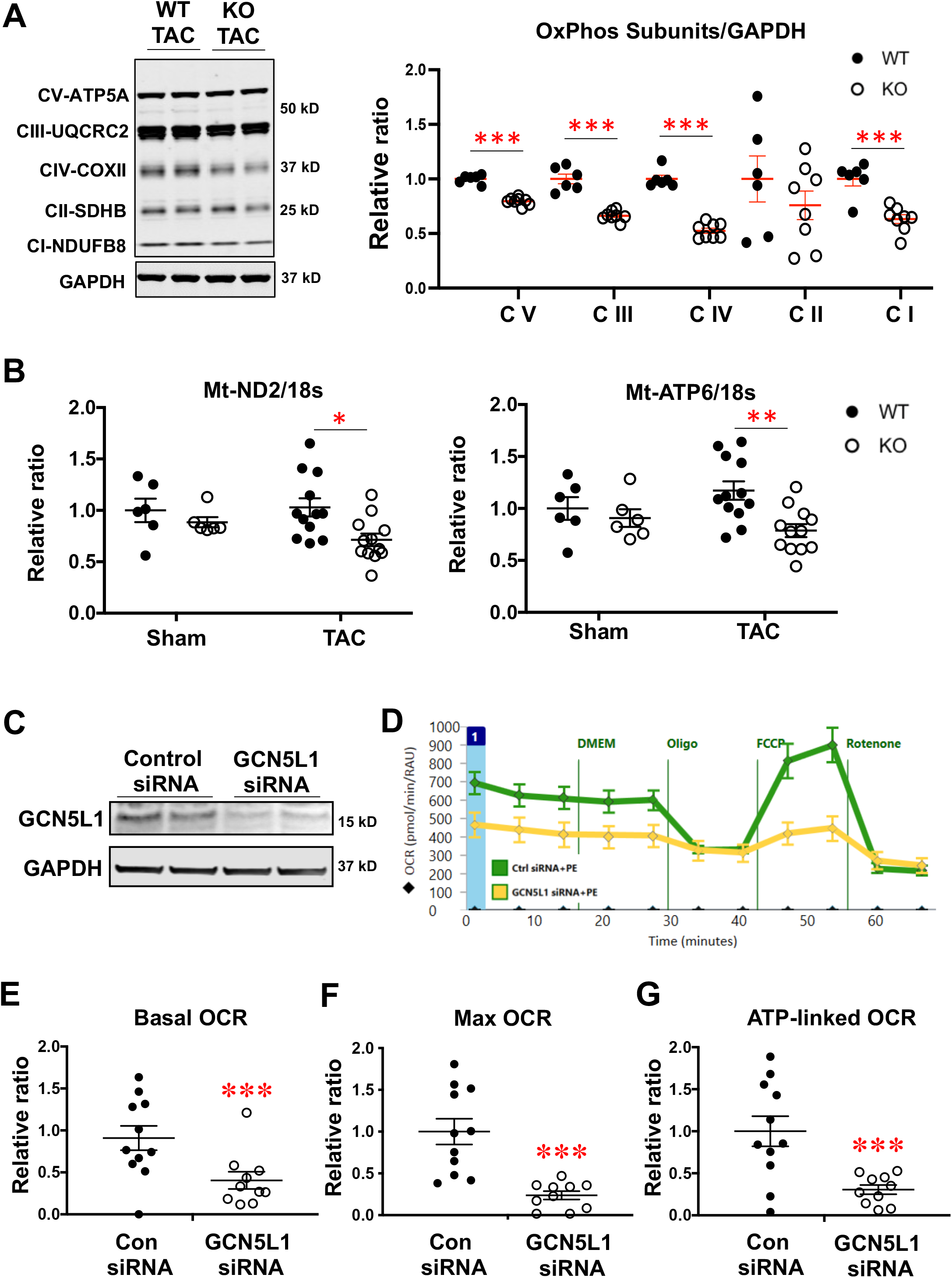
Loss of GCN5L1 expression in cardiomyocytes impairs mitochondrial bioenergetics. (A) OXPHOS cocktail western blot showed that mitochondrial proteins including NDUFB8 (complex I), SDHB (complex II), UQCRC2 (complex III), COXII (complex IV), and ATP5A (ATP synthase) were decreased in TAC cGCN5L1 KO mice, compared to WT controls (n=6 in WT, n=8 in KO). (B) Mitochondrial DNA copy number, assessed by mt-ND2/18s RNA and mt-ATP6/18s RNA, was reduced in cGCN5L1 KO mice subjected to TAC (n = 6 in sham groups and n = 12 in TAC groups). (C-G) Knockdown of GCN5L1 with siRNA in NRCMs resulted in impaired basal and maximum oxygen consumption rate (OCR), and decreased ATP synthesis-linked OCR, following treatment with phenylephrine (n = 8). Data are shown as means ± SEM. *P<0.05, **P<0.01, ***P<0.001.

As the decrease in ETC proteins and mtDNA levels suggested a negative impact on mitochondrial bioenergetics, we modelled how decreased GCN5L1 levels would affect bioenergetic output in neonatal rat cardiomyocytes (NRCMs). In NRCMs stimulated with the hypertrophy inducer phenylephrine, GCN5L1 siRNA knockdown resulted in greater cell surface area (Fig S3A) and fetal gene reprogramming (Fig S3B,C), matching the phenotype observed *in vivo* after TAC. Using Seahorse XF respirometry in control and hypertrophied NRCMs, we found that basal oxygen consumption rate (OCR), maximal OCR, and ATP synthesis-linked OCR were significantly decreased with GCN5L1 knockdown in the presence of phenylephrine (Fig 2C-G). Taken together, these findings suggest that GCN5L1 abundance governs mitochondrial bioenergetic output in cardiomyocytes during hemodynamic stress, and that reduced GCN5L1 levels promote energy deficits at the onset of hypertrophy.

### Adenoviral-mediated GCN5L1 overexpression restores bioenergetic output in hypertrophic cardiomyocytes

As loss of GCN5L1 expression negatively impacted mitochondrial bioenergetics in stressed hearts, we hypothesized that increasing GCN5L1 abundance may protect cardiomyocytes from hypertrophic remodeling. Indeed, adenoviral-mediated GCN5L1 overexpression (Fig 3A) inhibited fetal gene reprogramming in NRCMs treated with phenylephrine (Fig 3B,C), suggesting that increased GCN5L1 levels may be cardioprotective. We then examined whether GCN5L1 overexpression would reverse the bioenergetic decline observed in GCN5L1 depleted cells, and found that phenylephrine-treated NRVMs transduced with adenoviral GCN5L1 increased maximal OCR and ATP synthesis-linked OCR (Fig 3D-G). Combined, these data suggest that increased GCN5L1 abundance inhibits the progression of hypertrophic remodeling and limits bioenergetic dysfunction in stressed cardiomyocytes.

**Figure 3.**
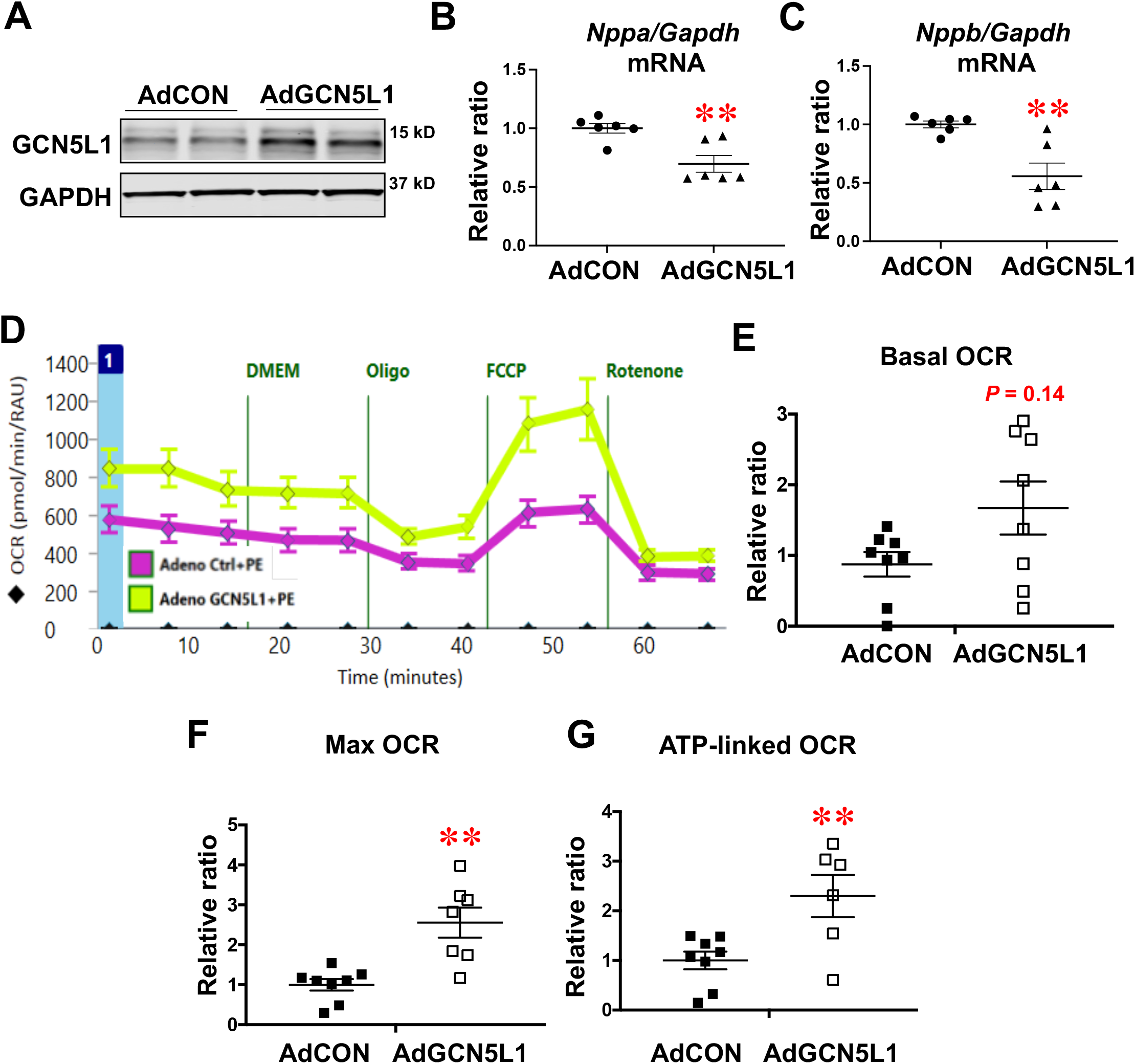
Adenoviral-mediated GCN5L1 overexpression inhibited cardiomyocyte hypertrophy and promoted mitochondrial bioenergetic output. (A-C) Adenoviral overexpression of GCN5L1 attenuated the PE induced cardiomyocyte hypertrophy, as measured by reduced fetal gene reprograming (n = 6). (D-G) Adenoviral overexpression of GCN5L1 in NRCMs increased maximal oxygen consumption rate (OCR) and ATP synthesis-linked OCR in the presence of PE (n = 8). Basal respiration was increased with GCN5L1 overexpression but did not reach statistical significance. Data are shown as means ± SEM. *P<0.05, **P<0.01, oooP<0.001.

### GCN5L1-mediated TFAM acetylation promotes increased mtDNA levels in cardiomyocytes

Finally, we investigated potential mechanisms underlying GCN5L1 control of mitochondrial bioenergetics in failing hearts. Consistent with previous reports (Manning et al., 2019; Thapa et al., 2020), we found that total mitochondrial protein acetylation levels were significantly decreased in cGCN5L1 KO mice relative to WT controls (Fig S4), supporting the concept that GCN5L1 is an important component of the mitochondrial acetylation regulatory machinery *in vivo*. As our initial studies suggested that a reduction in mtDNA levels may underpin respiratory deficits in cGCN5L1 KO mice, we examined whether the acetylation status of key regulators of mitochondrial gene expression was altered following cardiomyocyte-specific GCN5L1 depletion. We found that TFAM acetylation levels were decreased in cGCN5L1 KO mice hearts subjected to TAC relative to WT controls, which occurred in the absence of changes to TFAM protein levels (Fig 4A-C). To determine if GCN5L1 directly regulated TFAM acetylation levels *in vitro*, we transduced 293T cells with control or GCN5L1 adenovirus, and found that TFAM acetylation was significantly upregulated in response to GCN5L1 overexpression (Fig 4D).

**Figure 4.**
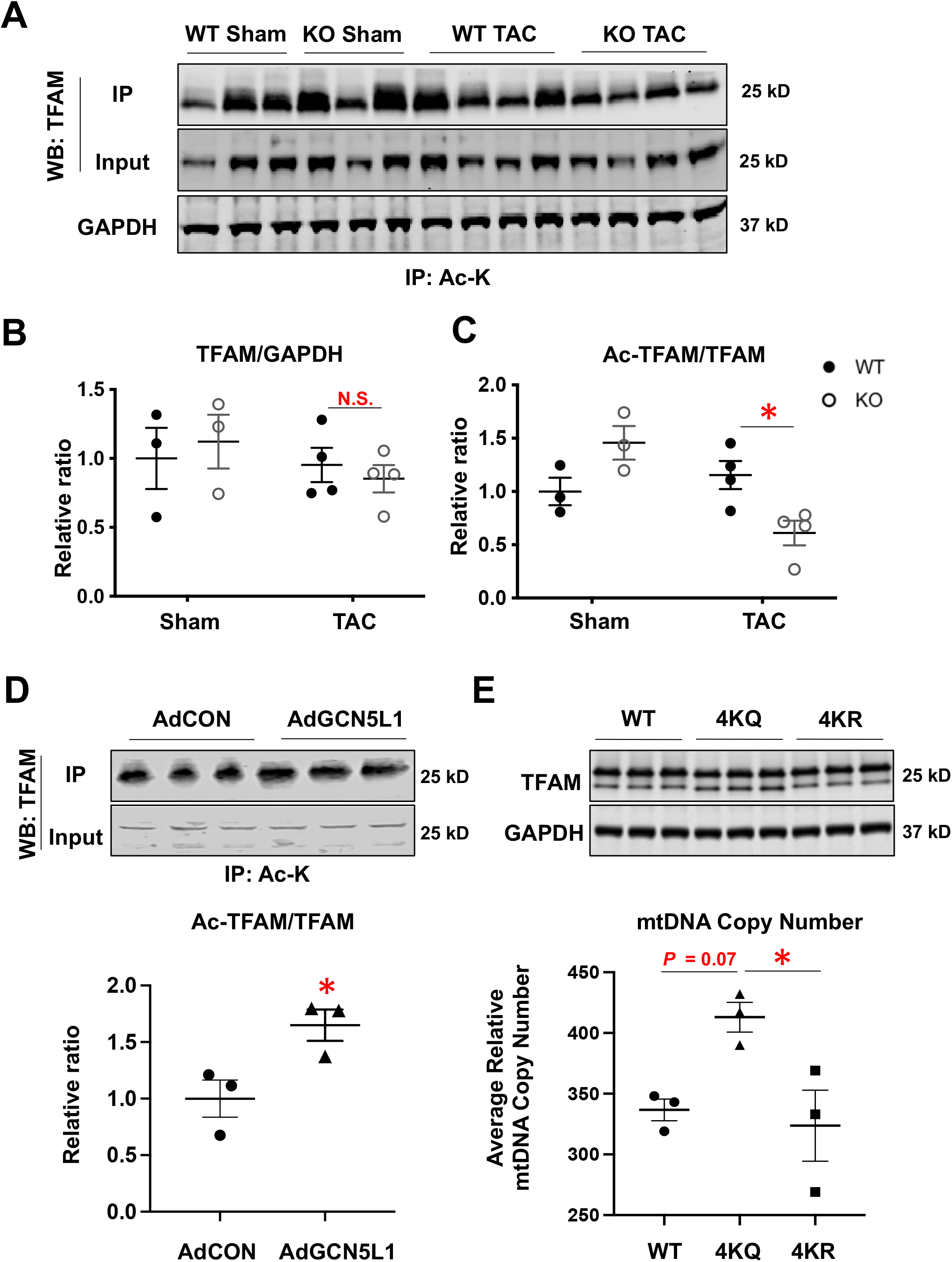
GCN5L1-mediated TFAM acetylation promotes increased mtDNA abundance. (A-C) The acetylation level of TFAM was decreased in cGCN5L1 KO mice subjected to TAC relative to WT controls, without changes in total TFAM protein abundance (n = 3-4). (D) Adenoviral overexpression of GCN5L1 in 293T cells increased the acetylation of TFAM (n = 3). (E) Overexpression of acetyl-mimetic 4KQ TFAM in AC16 cells increased total mtDNA copy number relative to the deacetylated-mimetic 4KR TFAM (n = 3). Data are shown as means ± SEM. P<0.05, **P<0.01, ***P<0.001.

Finally, to understand how acetylation of TFAM might impact mtDNA levels in cells, we generated expression plasmids for WT TFAM, and for two mutants where four lysine residues (K62/K76/K111/K118) were replaced with glutamine (4KQ) or arginine (4KR). These four lysine residues of TFAM have previously been reported to be acetylated in proteomic studies (Fang et al., 2020), and the substitutions made were designed to model the acetylated (4KQ) and deacetylated (4KR) state, respectively (see, e.g., Gorsky et al., 2016). Expression of the acetyl-mimetic 4KQ mutant in human cardiac AC16 cells led to a ~20% increase in mtDNA copy number relative to WT TFAM, which trended towards (but did not reach) statistical significance (*P* = 0.07). Conversely, expression of the deacetylated 4KR mimetic resulted in a significantly decreased mtDNA copy number relative to the acetylated 4KQ mimetic (Fig 4E), suggesting that acetylation of TFAM is correlated with an increase in mtDNA levels. Combined, these data suggest that GCN5L1 promotes bioenergetic output in stressed cardiomyocytes by maintaining mtDNA levels and mitochondrial respiratory function through the acetylation and activation of TFAM.

## Discussion

Lysine acetylation is recognized as a common post-translational modification of mitochondrial metabolic enzymes (Lu et al., 2009). In the heart, mitochondrial targets account for nearly 60% of total acetylated proteins and 64% of acetylation sites (Foster et al., 2013). The majority of acetylated proteins identified are enzymes involved in fatty acid metabolism, oxidative phosphorylation, or the tricarboxylic acid cycle (Foster et al., 2013), suggesting a possible regulatory role in mitochondrial energy metabolism. However, the role of global changes in lysine acetylation in the control of mitochondrial function remains controversial. For example, quantitative proteomic studies have found low acetylation occupancy rates of candidate regulatory lysine residues of metabolic enzymes, raising questions about the physiological impact of this post-translational modification (Baeza et al., 2016). More recently, a cardiac-focused study reported that global hyperacetylation of cardiac mitochondrial proteins, using a striated muscle-specific sirtuin 3 and carnitine acetyltransferase double knockout (SIRT3/CrAT DKO) mouse, had little negative impact on mitochondrial respiration or cardiac contractile function after TAC (Davidson et al., 2020). These findings suggested that non-specific global hyperacetylation of mitochondrial proteins is well tolerated under certain conditions, thereby questioning the potential physiological relevance of excess lysine acetylation in the hemodynamically stressed heart.

Intriguingly, rather than exacerbating the effects of hemodynamic stress as predicted, global mitochondrial protein hyperacetylation in SIRT3/CrAT DKO mice led to a 15-20% improvement in survivorship after TAC relative to WT mice over the course of the 16-week study, with a significant improvement being detected in the first three weeks after surgery (Davidson et al., 2020). These data suggest that rather than being inhibitory as first proposed, increased mitochondrial protein acetylation may actually promote some level of cardioprotection, particularly in the early stages of hemodynamic stress. Our new data presented here suggests that this may be related to the maintenance of mitochondrial bioenergetics, via GCN5L1-mediated TFAM acetylation, during the early hypertrophic response to pressure overload. In this context, a reduction in mitochondrial protein acetylation (via GCN5L1 downregulation or deletion) would limit mitochondrial bioenergetic output, while hyperacetylation (via SIRT3/CrAT deletion) may in fact be protective by maintaining TFAM acetylation and boosting cardiomyocyte energetics in the early stages of heart failure. Future studies, examining whether cardiac GCN5L1 downregulation remains deleterious in the context of SIRT3 deletion (where TFAM would presumably remain acetylated during TAC), will help to further delineate this pathway.

We previously identified several mitochondrial targets of GCN5L1, including fatty acid oxidation enzymes (long-chain acyl-CoA dehydrogenase [LCAD], short-chain acyl-CoA dehydrogenase [SCAD], and hydroxyacyl-CoA dehydrogenase [HADHA]) (Thapa et al., 2018; Thapa et al., 2017), electron transport chain (ETC) proteins (NDUFA9 from Complex I and ATP5a from ATP synthase) (Scott et al., 2012), and the glucose oxidation enzyme pyruvate dehydrogenase (PDHA1) (Thapa et al., 2019). Here we identified a new GCN5L1 target, TFAM, a transcription factor controlling mitochondrial DNA replication and mitochondrial metabolism. We found that the acetylation status of TFAM was reduced in cGCN5L1 KO mice hearts subjected to TAC in comparison with WT controls (Fig 4A). In 293T cells, overexpression of GCN5L1 increased the acetylation level of TFAM, further demonstrating that TFAM is likely a target of GCN5L1-mediated acetylation (Fig 4D). Indeed, the decreased mtDNA level observed in GCN5L1 KO mice hearts after TAC suggests that GCN5L1 potentially regulates mtDNA abundance through the acetylation of TFAM, and *in vitro* modeling of TFAM acetylation supports the hypothesis that hyperacetylation of this protein correlates with higher mtDNA levels (Fig 4E).

Proteomic detection of TFAM acetylation has been reported previously, with an initial characterization in a rat model demonstrating that TFAM was acetylated at a single residue in numerous organs (Dinardo et al., 2003). Several pieces of recent evidence have further suggested that TFAM acetylation is likely to have a regulatory effect on mtDNA gene expression and abundance. Using an acetyl-mimetic model, King et al. (2018) demonstrated that acetylated TFAM is less able to bind to DNA (via a reduced on-rate). This change may limit the ability of TFAM to maintain DNA compaction and allow fine-tuning of mtDNA-TFAM dynamics (King et al., 2018), leading to increased mtDNA transcription. Chemical acetylation of TFAM *in vitro* using acetyl-CoA led to several hyperacetylated lysine residues (including the four used in our mutant analyses), which may alter mtDNA topology (Fang et al., 2020). Finally, a contemporary study of TFAM post-translational modifications demonstrated that acetylated TFAM increased the processivity of the mitochondrial RNA polymerase POLRMT, allowing this enzyme to more easily move past TFAM roadblocks on mtDNA during transcription (Reardon and Mishanina, 2022).

In the context of the present study, these previous findings would suggest that GCN5L1-mediated acetylation of TFAM allows greater mitochondrial gene transcription and mtDNA abundance, which permits the maintenance of bioenergetic output to meet the demands of hemodynamic stress. We hypothesize that GCN5L1 downregulation in response to pressure overload (as occurs in human or mouse failing hearts; Fig 1A,B), or cardiomyocyte-specific deletion of GCN5L1 (Fig 1C-I), prevents the heart from upregulating bioenergetic output under stress, which aids the progression of cardiac dysfunction. Our findings suggest that mechanisms to maintain GCN5L1 abundance in the stressed heart may represent a potential new therapeutic approach in the treatment of heart failure.

## Acknowledgements

We thank the University of Pittsburgh Small Animal Ultrasonography Core, and funding for the echocardiography equipment from the NIH Instrumentation Program (NIH 1S10OD023684). The Center for Metabolism and Mitochondrial Medicine was supported by funding from the Pittsburgh Foundation to M.J.J. (MR2020 109502). The work was supported by NHLBI K08 HL157616, a UPMC HVI Fellows Research Award, and a Samuel and Emma Winters Foundation Award to M.Z.; by NHLBI K08 HL130604, American Heart Association Innovative Project Award #18IPA34170219, and R03 HL164393 to N.F.; and by NHLBI K22 HL116728, R01 HL132917, and R01 HL156874 to I.S.

## Author Contributions

M. Z., N.F. and I.S. designed the experiments. M.Z., N.F., D.T., M.S., J.M., C.M., X.Y., Z. P., and D.G. obtained the data. M.J.J. and M.N.S. provided critical reagents or expertise. M.Z., N. F., S.S., B.K., M.W.S. and I.S. analyzed the data. M.Z. wrote the manucript, I.S. edited the manuscript and made revisions.

## Inclusion and Diversity

We ensured sex balance in the experimental design by using equal numbers of male and female mice.

## STAR✶Methods

### Key Resources

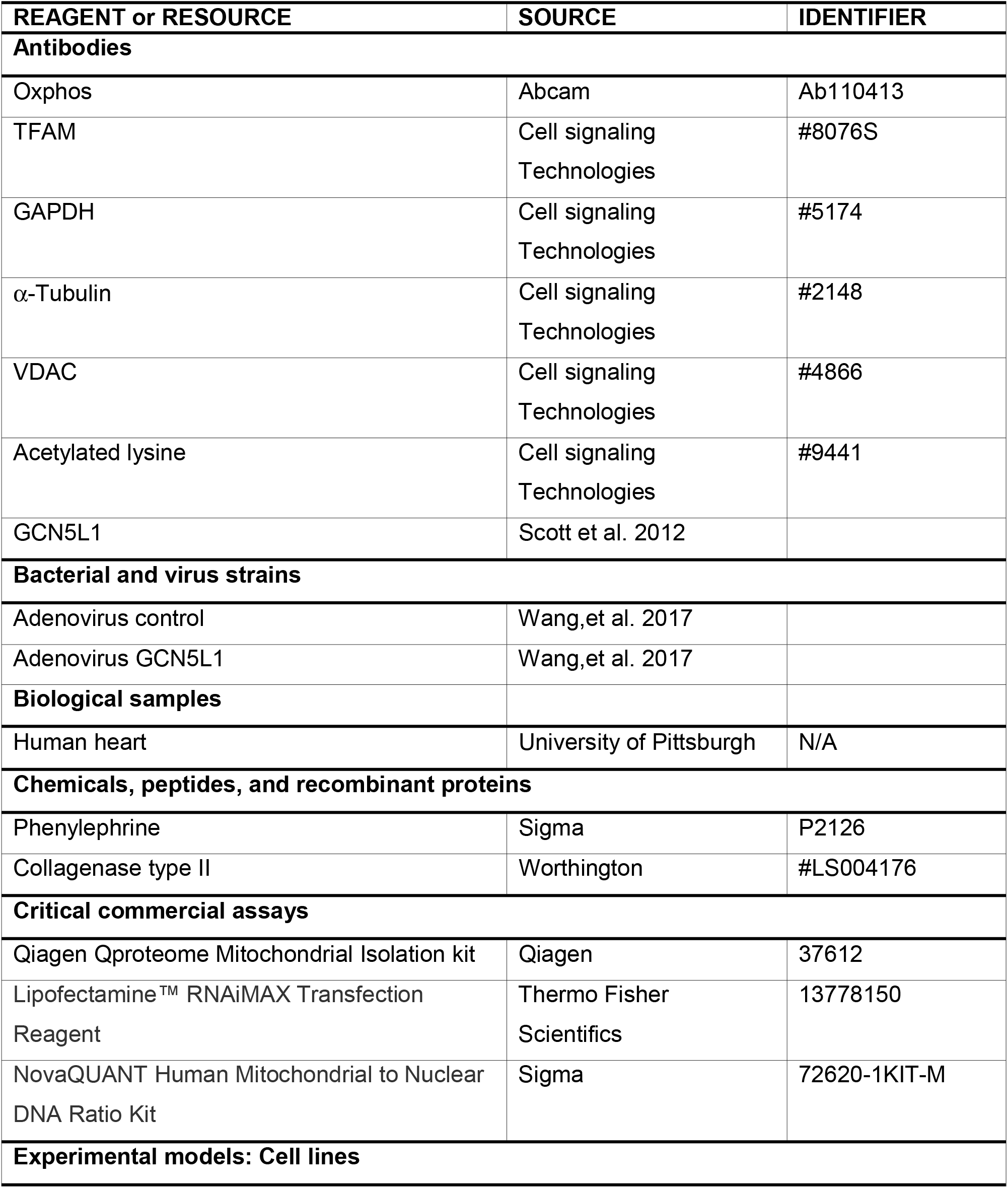
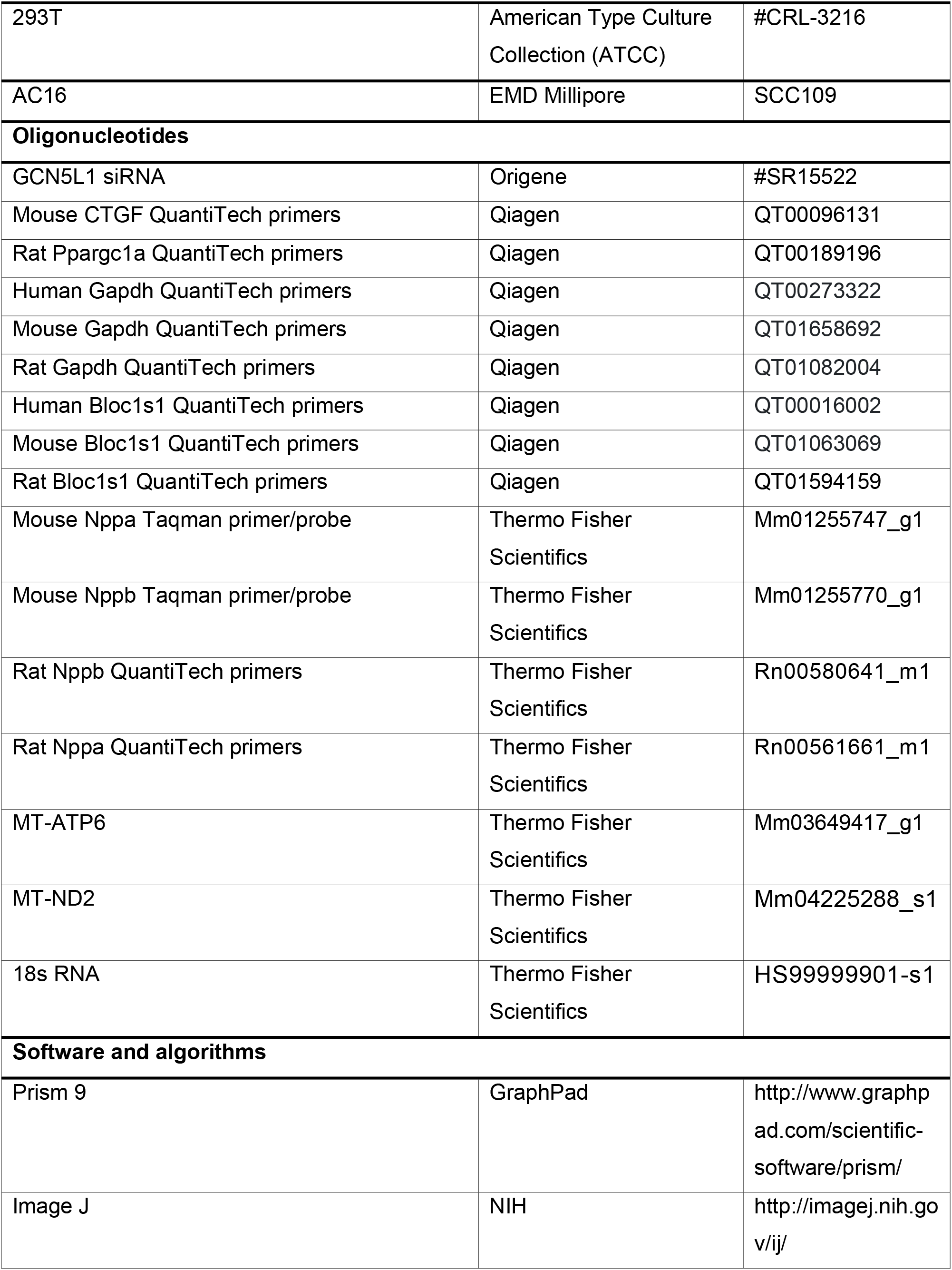

| REAGENT or RESOURCE | SOURCE | IDENTIFIER |
| --- | --- | --- |
| **Antibodies** |  |  |
| Oxphos | Abcam | Ab110413 |
| TFAM | Cell signaling Technologies | #8076S |
| GAPDH | Cell signaling Technologies | #5174 |
| α-Tubulin | Cell signaling Technologies | #2148 |
| VDAC | Cell signaling Technologies | #4866 |
| Acetylated lysine | Cell signaling Technologies | #9441 |
| GCN5L1 | Scott et al. 2012 |  |
| **Bacterial and virus strains** |  |  |
| Adenovirus control | Wang,et al. 2017 |  |
| Adenovirus GCN5L1 | Wang,et al. 2017 |  |
| **Biological samples** |  |  |
| Human heart | University of Pittsburgh | N/A |
| **Chemicals, peptides, and recombinant proteins** |  |  |
| Phenylephrine | Sigma | P2126 |
| Collagenase type II | Worthington | #LS004176 |
| **Critical commercial assays** |  |  |
| Qiagen Qproteome Mitochondrial Isolation kit | Qiagen | 37612 |
| Lipofectamine™ RNAiMAX Transfection Reagent | Thermo Fisher Scientifics | 13778150 |
| NovaQUANT Human Mitochondrial to Nuclear DNA Ratio Kit | Sigma | 72620-1KIT-M |
| **Experimental models: Cell lines** |  |  |
| 293T | American Type Culture Collection (ATCC) | #CRL-3216 |
| AC16 | EMD Millipore | SCC109 |
| **Oligonucleotides** |  |  |
| GCN5L1 siRNA | Origene | #SR15522 |
| Mouse CTGF QuantiTech primers | Qiagen | QT00096131 |
| Rat Ppargc1a QuantiTech primers | Qiagen | QT00189196 |
| Human Gapdh QuantiTech primers | Qiagen | QT00273322 |
| Mouse Gapdh QuantiTech primers | Qiagen | QT01658692 |
| Rat Gapdh QuantiTech primers | Qiagen | QT01082004 |
| Human Bloc1s1 QuantiTech primers | Qiagen | QT00016002 |
| Mouse Bloc1s1 QuantiTech primers | Qiagen | QT01063069 |
| Rat Bloc1s1 QuantiTech primers | Qiagen | QT01594159 |
| Mouse Nppa Taqman primer/probe | Thermo Fisher Scientifics | Mm01255747_g1 |
| Mouse Nppb Taqman primer/probe | Thermo Fisher Scientifics | Mm01255770_g1 |
| Rat Nppb QuantiTech primers | Thermo Fisher Scientifics | Rn00580641_m1 |
| Rat Nppa QuantiTech primers | Thermo Fisher Scientifics | Rn00561661_m1 |
| MT-ATP6 | Thermo Fisher Scientifics | Mm03649417_g1 |
| MT-ND2 | Thermo Fisher Scientifics | Mm04225288_s1 |
| 18s RNA | Thermo Fisher Scientifics | HS99999901-s1 |
| **Software and algorithms** |  |  |
| Prism 9 | GraphPad | http://www.graphpad.com/scientific-software/prism/ |
| Image J | NIH | http://imagej.nih.gov/ij/ |

### Resource Availability

#### Lead contact

Further information, or requests for reagents or resources, should be directed to the lead contact, Dr. Iain Scott.

#### Materials availability

All unique/stable reagents generated in this study are available from the lead contact with a completed Materials Transfer Agreement.

#### Experimental model and subject details

Study procedures were approved by the University of Pittsburgh Animal Care and Use Committees (IACUC) in accordance with National Institutes of Health guidelines. Floxed GCN5L1 KO mice were reported previously (Manning et al., 2019). Cardiac-specific GCN5L1 KO mice (cGCN5L1 KO mice) were generated by crossing the floxed GCN5L1 mice with αMHC-Cre mice (Jackson Lab, Stock # 011038).

Human heart failure tissues were obtained from transplant recepients with non-ischemic cardiomyopathy at the University of Pittsburgh Medical Center with Institutional review board (IRB) approval and informed consents. The control human heart tissues were obtained from donor hearts which did not meet the criteria for transplant with IRB approval and informed consents. All samples were preserved in cold cardioplegic solution and transported to the laboratory. Heart tissue was then snap frozen in liquid nitrogen and then stored at −80°C until assay.

### Detailed Methods

#### TAC procedure

TAC procedure was performed as previously described (Yang et al., 2022). Briefly, sterile surgery was performed in our dedicated facility. Anesthesia was induced with 3.0% isoflurane, and maintained by ventilator at 2.0% isoflurane. Skin was shaved and steriled with povidone-iodine. An incision was made to the immediate left of the sternum, the 2^nd^ and 3^rd^ ribs were cut, and blunt isolation was done to expose the aortric arch. A 7-0 proline suture tie was placed around the aortic arch between the bronchiocephalic artery and left common carotid artery, and tightened around a 27- or 26-gauge needle. Once the tie was fixed, the needle was removed, and the suture/stenosis remained. The chest was then closed with a 6-0 proline suture and 5-0 silk suture. Serial echocardiography was performed in sedated mice (TAC) and conscious mice (12-months-old) as described previously (Yang et al., 2022).

#### Mitochondrial DNA copy number and mitochondrial protein level assessment

Mitochondrial DNA and genomic DNA were isolated from mouse hearts using QIAamp Fast DNA Tissue Kit (Qiagen #51404) according to the manufacturer’s instructions. mtDNA copy number was assessed with Mitochondrially Encoded NADH:Ubiquinone Oxidoreductase Core Subunit 2, (mt-ND2), or Mitochondrially Encoded ATP Synthase Membrane Subunit 6, (mt-ATP6)/genomic DNA (18s RNA) ratio as previously described (Jiang et al., 2017). The Taqman primers were obtained from Thermo Fisher Scientific (MT-ATP6: Mm03649417_g1, MT-ND2: Mm04225288_s1, 18 sRNA gene: Hs99999901_s1). Total mitochondrial DNA copy number was analyzed using the NovaQUANT Human Mitochondrial to Nuclear DNA Ratio Kit (Sigma #72620-1KIT-M). Mitochondrial proteins level was evaluated by Western Blot using OXPHOS cocktail antibodies (Abcam, #ab110413). Mitochondrial proteins were isolated using the Qiagen QProteome Mitochondrial Isolation kit according to the manufacturer’s instruction (Thapa et al., 2019).

#### Cardiac myocytes and cell line studies

Neonatal rat cardiomyocytes (NRCMs) were freshly isolated as described previously (Zhang et al., 2020). Briefly, hearts were quickly removed from one to three day-old Sprague Dawley neonates, and cardiac myocyte isolation was achieved by digestion with 0.04% trypsin and type II collagenase (0.4mg/ml; Worthington #LS004176) in Krebs-Henseleit bicarbonate buffer at 37°C. Non-cardiomyocyte cells were removed by rapid attachment (90 minutes incubation in culture dishes). Cardiomyocytes were plated at the density of 2×10^5^/ml in DMEM containing 10% FBS and 0.1mM BrdU to prevent the growth of fibroblasts. Twenty-four hours after cells plating, the medium was changed to serum-free DMEM containing 0.1% insulin-transferrin-selenium (Thermo Fisher Scientific #41400045). 293T cells were obtained from ATCC (#CRL-3216), and cultured in DMEM containing 10% FBS. AC16 cells were obtained from EMD Millipore (#SCC109) and maintained in DMEM containing 10% FBS.

#### Mitochondrial bioenergetics measurements in cultured cardiac myocytes

Oxygen consumption rate (OCR) was measured in NRCMs transfected with control siRNA, GCN5L1 siRNA, control adenovirus, or GCN5L1 adenovirus using the Seahorse XF system (Seahorse Bioscience) as described previously (Thapa et al., 2019). Basal OCR in each well was measured, followed by serial treatment with FCCP (7.5 μmol/L) and rotenone (2 μmol/L). Each experiment was repeated to ensure reproducibility, and the data presented are technical replicates (N = 8-11) of a single representative study. NRCMs were transfected with GCN5L1 siRNA (Origene # SR15522) and control siRNA using Lipofectamine™ RNAiMAX Transfection Reagent (Thermo Fisher Scientific #13778150) according to the manufacturer’s protocol. The adenovirus overexpressing GCN5L1 was previously reported (Wang et al., 2017). Briefly, adenoviruses were produced using the Adeasy Adenoviral system (Agilent), and a multiplicity of infection (MOI) of 10 was used for all the experiments. NRCMs were treated with phenylephrine (20 μM; Sigma #P6126), 24 hours after siRNA transfection, for 48 hours before use or harvest. The human TFAM WT, 4KQ (K62Q/K76Q/K111Q/K118Q), and 4KR (K62R/K76R/K111R/K118R) plasmids were custom synthesized by Genscript, and transiently transfected for 48 hours into AC16 cells using Lipofectamine as above.

#### Western blot and immunoprecipation

Protein extracts were prepared in RIPA lysis buffer (Thermo Fisher Scientific #89900) from snap-frozen heart tissues. Protein concentration was measured by BCA assay (Thermo Fisher Scientific #23227). Immunoprecipitation of tissue/cell lysates was performed in RIPA buffer, with equal amounts of protein used for immunoprecipitation assay with rabbit acetyl-lysine (AcK) antibody (Cell Signaling Technology #9441) and protein G agarose beads (Cell Signaling Technology #37478) overnight at 4°C. Beads were washed 4 times with RIPA buffer, and then eluted directly with LDS sample buffer (Thermo Fisher Scientific #B0007) at 95°C for 5 mins. Protein electrophoresis was performed on 4–12% Bis-Tris NuPage gels (Thermo Fisher Scientific #NW04120BOX). The Bio-Rad Trans-Blot Turbo Transfer System was used for protein transfer to nitrocellulose membranes. The related secondary antibodies used were from LI-COR Biosciences, and blots were quantified using Image J software (NIH). GCN5L1 antibody was used as previously reported (Scott et al., 2012). Oxphos Cocktail antibody (#ab110413) was from Abcam. The TFAM (#8076S), α-tubulin (#2148), GAPDH (#5174), and VDAC (#4866) antibodies were from Cell Signaling Technologies.

#### Quantitative RT-PCR

Total RNA was extracted from NRCMs or snap-frozen heart tissues using TRIzol reagent (Thermo Fisher Scientific). Reverse transcription was conducted using the High-Capacity cDNA Reverse Transcription Kit (Thermo Fisher Scientific). Taqman primers (Thermo Fisher Scientific) were used for quantitative RT-PCR analysis: mouse PPARGC1A (Mm01208835_m1), and mouse TFAM (Mm00447485_m1).

#### Statistics

Data were compared within groups using one-way or two-way ANOVA with Tukey’s Post Hoc Test using GraphPad Prism version 9.0. Unpaired student’s *t*-test was used for comparison between two groups. All tests were two-tailed and a *P* value of less than 0.05 was considered significant. All values are represented as the mean ± SEM.

---

**Figure S1:**
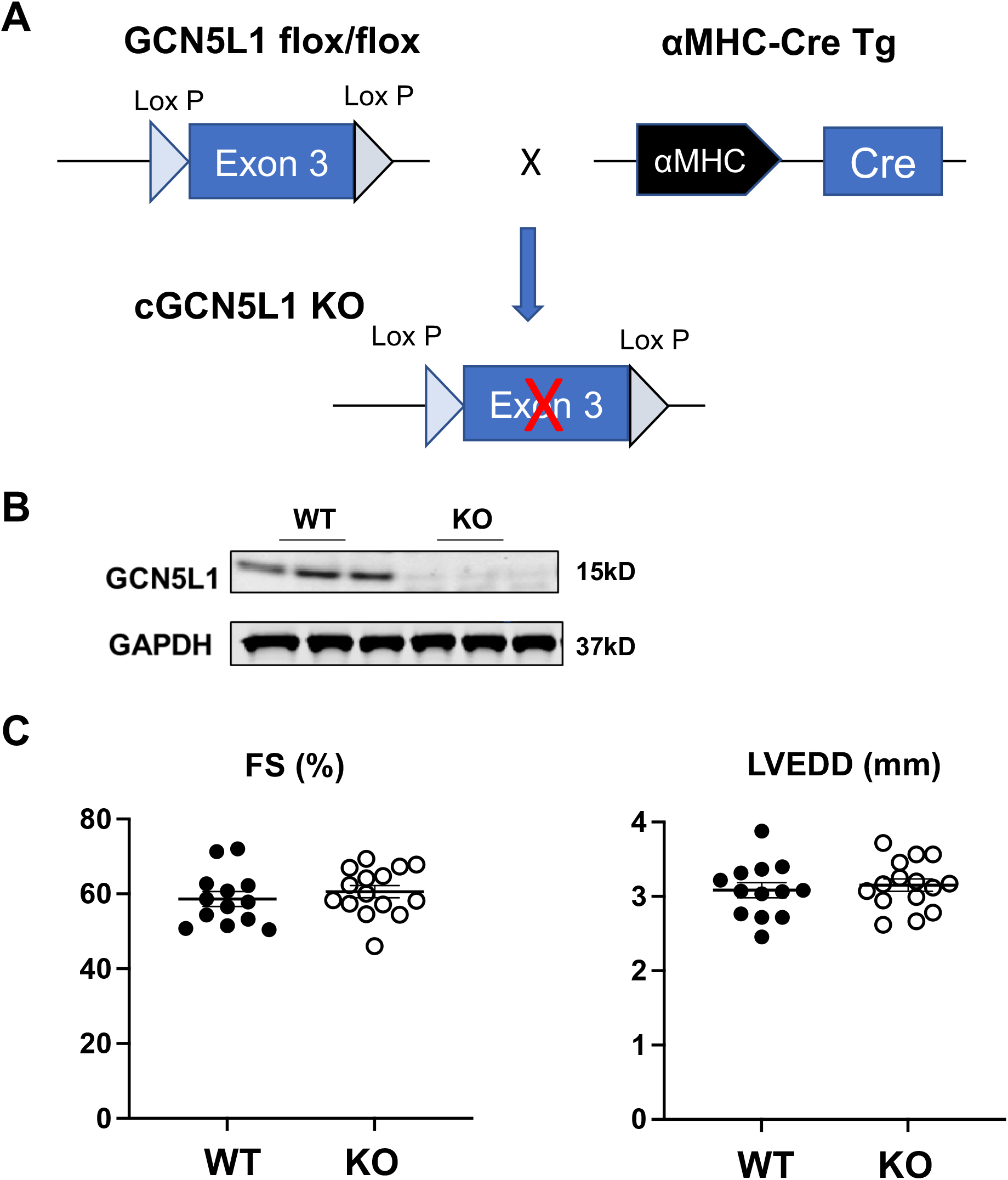
Generation of cardiac-specific GCN5L1 KO mice. (A)Construction of GCN5L1 conditional knockout. (B) Validation of cGCN5L1 KO deletion efficiency in the heart. (C) Cardiac function and chamber size of cGCN5L1 KO mice was preserved at one year old (n = 13-15).

**Figure S2:**
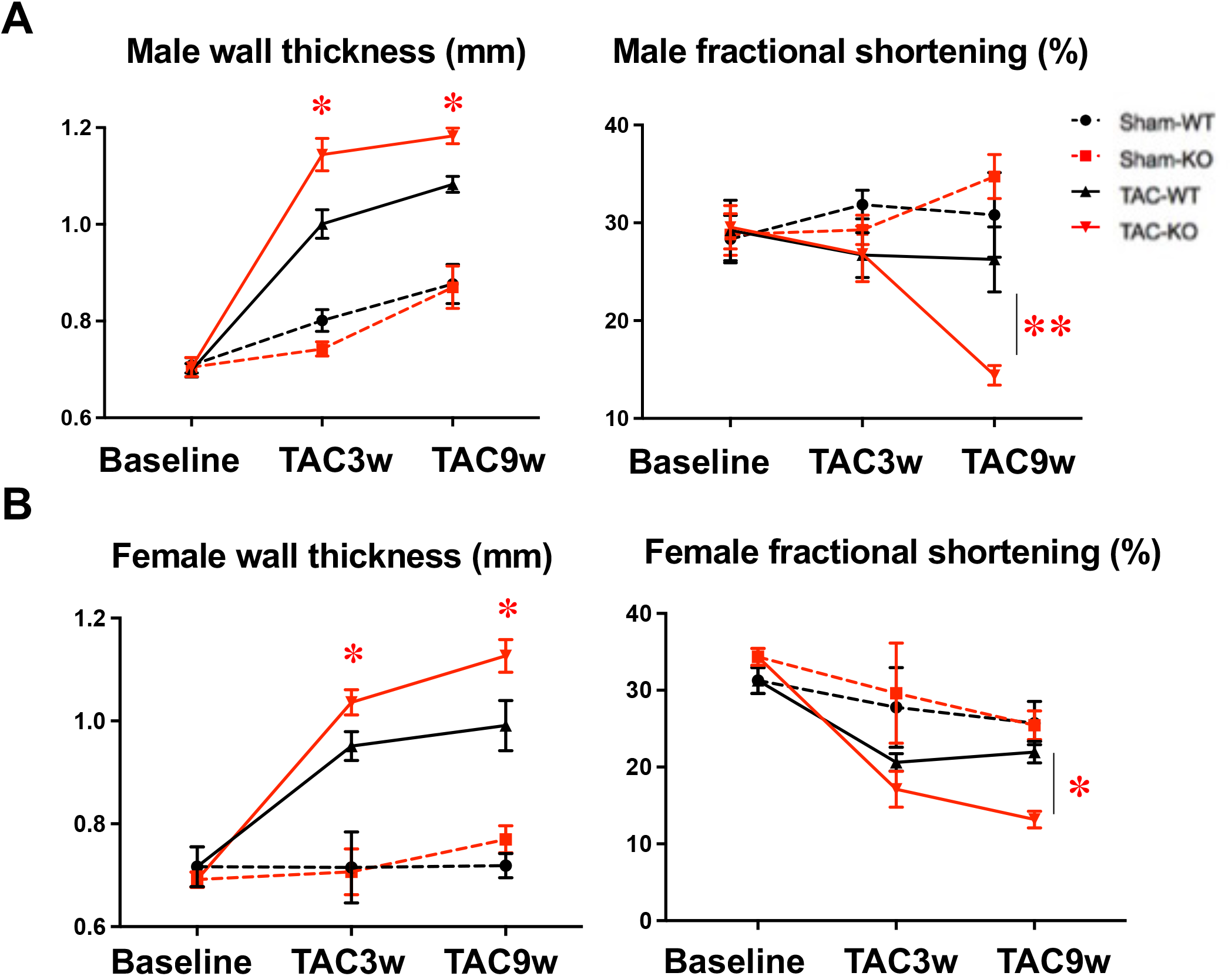
Both male and female cGCN5L1 KO mice display exacerbated heart failure progression after TAC. (A) Male cGCN5L1 KO mice showed increased wall thickness and decreased fractional shortening in comparison to WT controls after TAC (n=7). (B) Female cGCN5L1 KO mice displayed a similar response to TAC as male mice (n=8). *P<0.05 vs. TAC-WT group

**Figure S3:**
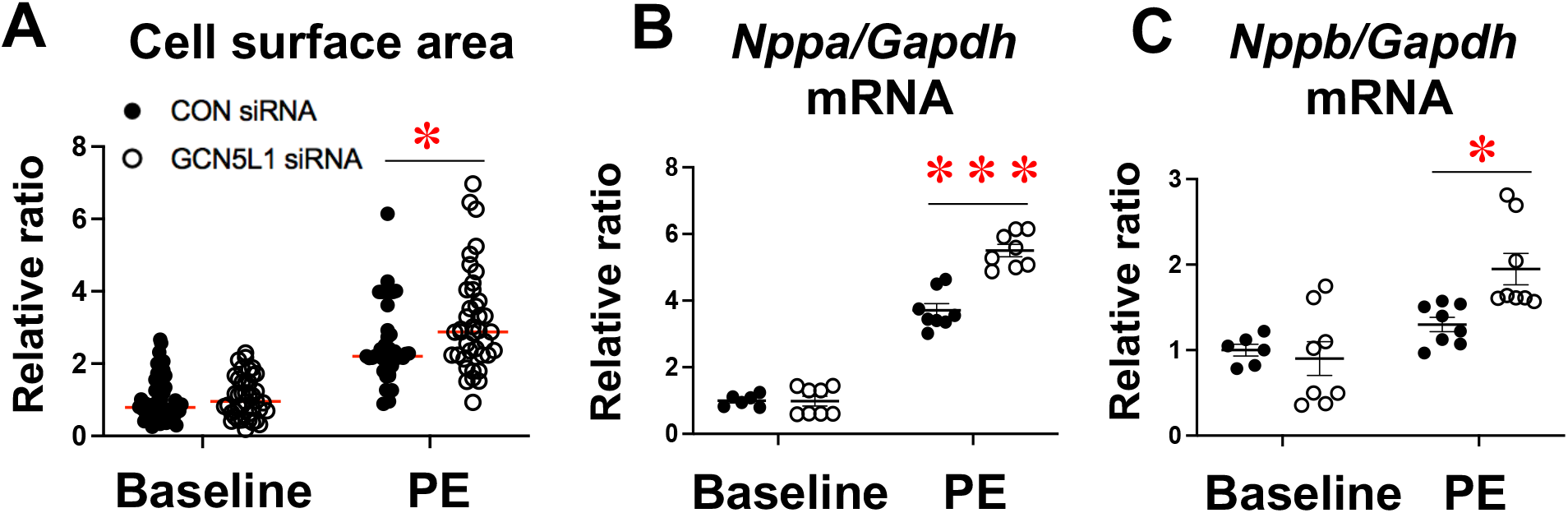
GCN5L1 knockdown in NRCMs results in cellular hypertrophy. siRNA-mediated depletion of GCN5L1 in neonatal rat cardiomyocytes results in increased cellular surface area and upregulated natriuretic peptide gene expression. (N = 6-50, * = *P* < 0.05; *** = *P* < 0.001)

**Figure S4:**
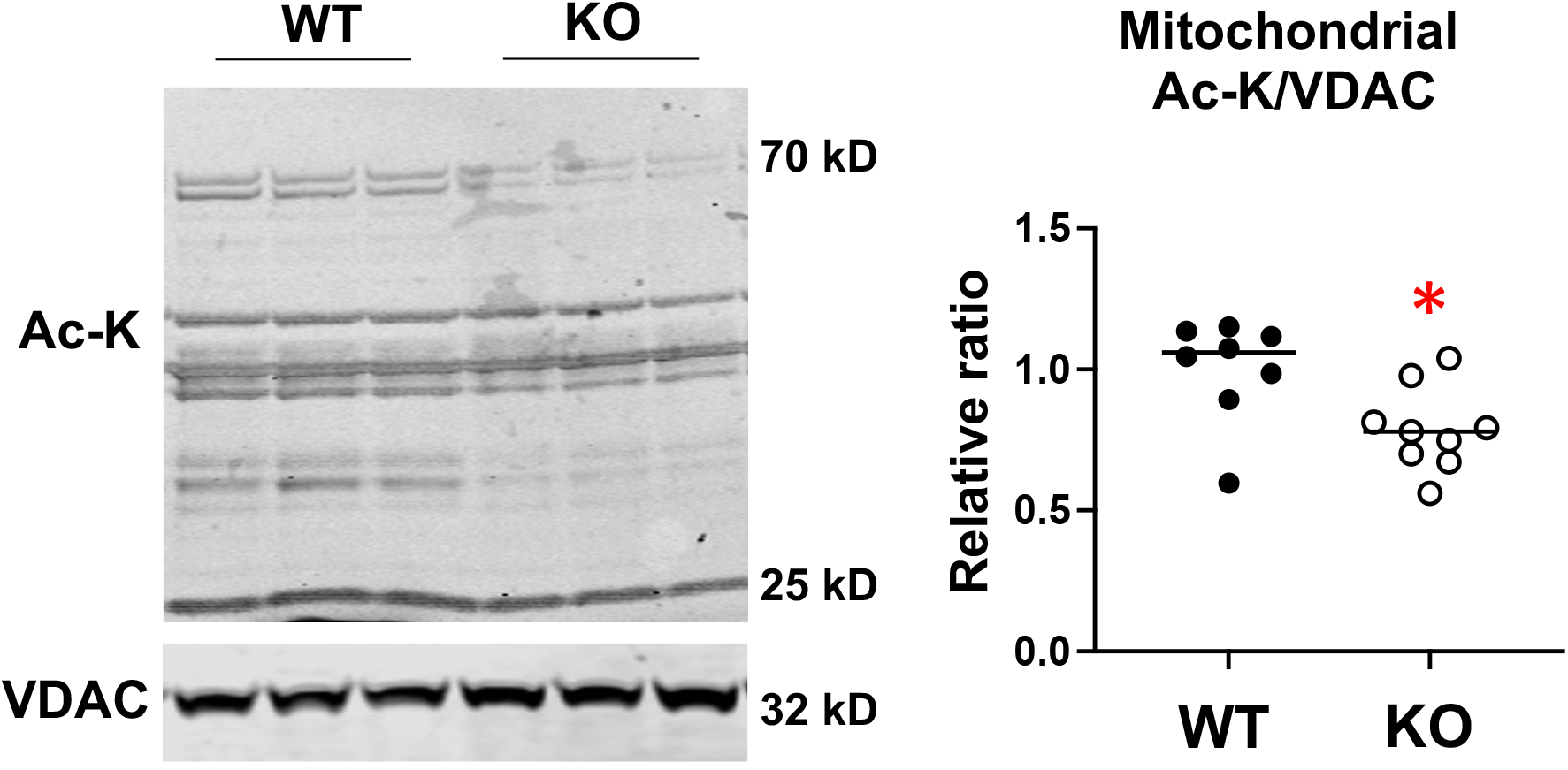
Reduced mitochondrial protein acetylation in GCN5L1 KO mice. Cardiomyocyte-specific deletion of GCN5L1 resulted in a significant decrease in mitochondrial protein lysine acetylation at baseline. (N = 8-9, * = *P* < 0.05).

